# Assessing the role of inhibition in stabilizing neocortical networks requires large-scale perturbation of the inhibitory population

**DOI:** 10.1101/123950

**Authors:** Sadra Sadeh, R. Angus Silver, Thomas Mrsic-Flogel, Dylan Richard Muir

## Abstract

Neurons within cortical microcircuits are interconnected with recurrent excitatory synaptic connections that are thought to amplify signals (Douglas and Martin, 2007), form selective subnetworks (Ko et al., 2011) and aid feature discrimination. Strong inhibition (Haider et al., 2013) counterbalances excitation, enabling sensory features to be sharpened and represented by sparse codes (Willmore et al., 2011). This “balance” between excitation and inhibition makes it difficult to assess the strength, or gain, of recurrent excitatory connections within cortical networks, which is key to understanding their operational regime and the computations they perform. Networks of neurons combining an unstable high-gain excitatory population with stabilizing inhibitory feedback are known as inhibition-stabilized networks (ISNs; Tsodyks et al. 1997). Theoretical studies using reduced network models predict that ISNs produce paradoxical responses to perturbation, but experimental perturbations failed to find evidence for ISNs in cortex (Atallah et al., 2012). We re-examined this question by investigating how cortical network models consisting of many neurons behave following perturbations, and found that results obtained from reduced network models fail to predict responses to perturbations in more realistic networks. Our models predict that a large proportion of the inhibitory network must be perturbed in order to robustly detect an ISN regime in cortex. We propose that wide-field optogenetic suppression of inhibition under a promoter targeting all inhibitory neurons may provide a perturbation of sufficient strength to reveal the operating regime of cortex. Our results suggest that detailed computational models of optogenetic perturbations are necessary to interpret the results of experimental paradigms.

**Significance statement:** Many useful computational mechanisms proposed for cortex require local excitatory recurrence to be very strong, such that local inhibitory feedback is necessary to avoid epileptiform runaway activity (an “inhibition-stabilized network” or “ISN” regime). However, recent experimental results suggest this regime may not exist in cortex. We simulated activity perturbations in cortical networks of increasing realism, and found that in order to detect ISN-like properties in cortex, large proportions of the inhibitory population must be perturbed. Current experimental methods for inhibitory perturbation are unlikely to satisfy this requirement, implying that existing experimental observations are inconclusive about the computational regime of cortex. Our results suggest that new experimental designs, targetting a majority of inhibitory neurons, may be able to resolve this question.

## Introduction

Inspired by experimental observations of a repeated, “canonical” architecture for cortex (Creutzfeldt, 1977; Rockel et al., 1980; Muir et al., 2011), several authors have proposed that a concomitant canonical function might also exist — a fundamental computational basis, common to all cortical areas (e.g. Szentagothai, 1978; Douglas et al., 1989). How can this computational principle be discovered? A frequently-applied approach in reverse-engineering a complex dynamical system is to measure the response of a system to a perturbing stimulus. This technique has been applied to cortex in the past (Douglas et al., 1989), but recent methodological advances permit targeted stimulation or suppression of chosen neuronal populations, through genetic targetting of light-sensitive ion channels and pumps (*optogenetics*; Boyden et al. 2005; Atallah et al. 2012; Han and Boyden 2007; Zhang et al. 2007). Optogenetic stimulation can be used to drive or suppress the activity of genetically-defined cell classes, or cortical populations with particular projection targets. This approach confers the possibility to use carefully targeted perturbations to observe and detect the computational mode of cortex. However, due to the prevalence of recurrent interactions in cortical networks, the outcome of such a perturbation may be unintuitive or difficult to predict. For this reason, computational modelling of perturbations is required to relate network architectures and operating regimes to the expected result of a particular perturbation, and to guide the choice of an appropriate experimental perturbation to optimally test hypotheses. Here we take as specific example the question of quantifying the excitatory / inhibitory balance in cortex, with a particular focus on mouse visual cortex.

Network computational mechanisms that rely on recurrent processing of information within cortex can be flexible and powerful (Hopfield, 1982; Douglas and Martin, 2007; Hopfield, 2015). Many computational models for mammalian cortex require strong recurrent excitation, which consequently must be balanced by strong local inhibition to maintain stability of the cortical network (Hahnloser, 1998; Rutishauser and Douglas, 2009; Neftci et al., 2013; Muir and Cook, 2014). Networks with this property are known as *inhibition-stabilized networks*, or ISNs (Tsodyks et al., 1997; Ozeki et al., 2009; Litwin-Kumar et al., 2016). An alternative configuration of cortical networks could rely on a weak excitatory population that is intrinsically stable, which would support different computational mechanisms not relying on strong excitatory recurrence. The question of which balanced regime mammalian neocortex operates in is therefore of experimental interest, as this constrains the type of computations that could be supported by cortex. Anatomical and physiological estimates suggest that recurrent excitation is very strong, especially in the superficial layers of cortex (Binzegger et al., 2004; Lefort et al., 2009). Similarly, observations of epileptiform activity when inhibition is blocked in cortex suggest that inhibitory feedback is required for stability of the cortical network (Avoli et al., 1995; Mann et al., 2009). However an ISN regime may also be detected functionally by experimentally perturbing the dynamics of cortical activity and observing the response of the network.

Here we analyse theoretical and simulation models of cortical networks, to determine the conditions under which an inhibitory perturbation evokes a measurable paradoxical response in the network, which can be used to infer the computational regime of cortex (Tsodyks et al., 1997). We then examine whether existing methods for perturbation of cortical activity — for example, electrical stimulation by injecting currents into inhibitory neurons; perfusion of the brain with chemical agonists or antagonists of inhibitory synaptic receptors (Bowery et al., 1984); or optogenetics — will be able to reveal evidence for an ISN regime in cortex.

## Results

Simple ISNs display counterintuitive dynamics when inhibitory activity is perturbed by increasing or decreasing excitatory input into inhibitory neurons. If inhibition is reduced by removing input then the network effect is to *increase* the activity of inhibitory neurons; conversely, if extra input is provided to inhibitory neurons then the network responds by *decreasing* their activity (Fig. 1). This has been termed the “paradoxical” inhibitory response (Tsodyks et al., 1997), and arises through nonlinear network dynamics introduced by unstable excitatory feedback. This counterintuitive effect of perturbing inhibition has been put forward as a signature of ISN dynamics that could be detected in cortical networks (Tsodyks et al., 1997). This is an experimentally accessible metric, since neurons are often being recorded and activated at the same time. When the entire inhibitory population of an ISN is perturbed simultaneously, then the paradoxical effect emerges as in Fig. 1. However, under typical experimental conditions only a fraction of the inhibitory population can be perturbed. This raises the question of whether the paradoxical effect will be observed if only portions of the inhibitory population are perturbed. Recent results based on direct activation and suppression of the inhibitory network (Atallah et al., 2012) did not reveal evidence for a paradoxical inhibitory response. Based on these results, some authors have inferred that an ISN regime may not exist in the superficial layers of mouse visual cortex (Litwin-Kumar et al., 2016). It remains unclear whether experimental methods for perturbing inhibition will be sufficient to reveal a signature of ISN dynamics.

**Figure 1:**
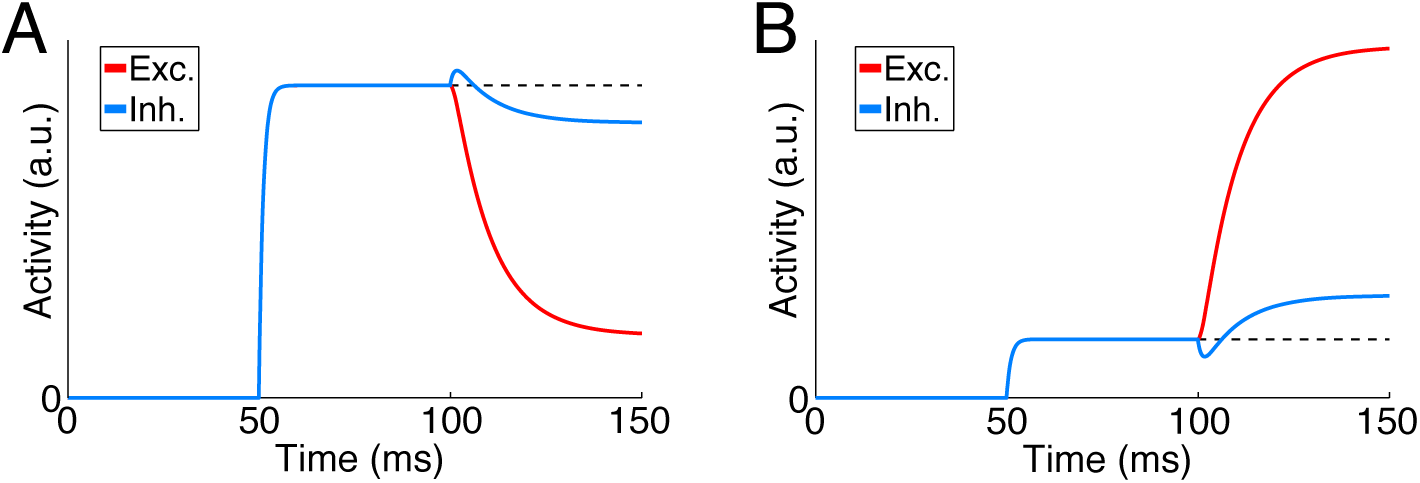
Globally perturbing the inhibitory network gives rise to a paradoxical inhibitory response, in inhibition-stabilized networks. The effect on activity of excitatory and inhibitory neurons, of increasing (A) and decreasing (B) input to the inhibitory population. At 50 ms input is injected to all neurons. At 100 ms the input to the inhibitory population only is perturbed. Note that increasing inhibitory input results in a counterintuitive decrease in overall inhibitory activity, and vice versa. Parameters: {*w_E_, w_I_, τ*} = {5, 20, 10 ms}. Dashed line for reference with pre-perturbation activity.

## Perturbations of homogeneous networks of firing-rate neurons in ISN and non-ISN regimes

To explore the properties of ISNs and non-ISNs and investigate how they respond to perturbations over a wide range of parameters, we first developed a simple analytically tractable model of a cortical network. For this we used nonspiking linear-threshold neuron models, since they provide a good approximation to the F-I curves of adapted cortical neurons (Ermentrout, 1998). Net-works were built using homogeneous synaptic connectivity and equal numbers of excitatory and inhibitory neurons (see Materials and Methods). In these models we simulated synaptic inputs by injecting currents proportional to pre-synaptic activity.

We analytically examined the stability and dynamic properties of this network, to determine the conditions under which it operates in an ISN regime. The stability of networks was determined by expressing all synaptic connections between pairs of neurons as a weight matrix *W*, and then analysing the properties of this matrix. Each network has an associated property known as the *largest real eigenvalue* λ_1_, which depends on the strength of excitation and inhibition within the network, and the dynamical properties of the network (see Materials and Methods). If this value is large (i.e. *λ*_1_ > 1), then the network can become unstable; this is because a pattern of activity in the network can become amplified through local recurrent feedback, and the firing activity of the neurons involved could increase without bound. Alternatively, if λ_1_ ≤ 1 then the activity of all neurons in the network is guaranteed not to increase without bound and this is defined as a *stable* network.

For a network to operate in an ISN regime, the network must be unstable in the absence of inhibition, yet stable with inhibitory feedback (Tsodyks et al.,1997). By setting the synaptic strength of inhibition *w_I_* to zero, we found that the excitatory network is unstable (i.e. λ_1_ > 1) when the total synaptic strength contributed by a single excitatory neuron (*w_E_*) is greater than 1. The interpretation of *w_E_* > 1 is that on average in an active network, a single spike from an excitatory neuron leads to at least one extra spike in the rest of the network.

To ensure stability in the entire network (i.e. λ_1_ ≤ 1 in the presence of inhibitory feedback), we found a constraint relating the strength of excitation and inhibition that guarantees local inhibition is strong enough to keep recurrent excitation in check. For networks operating in the ISN regime, the relative strengths of excitation and inhibition must satisfy 1 < *w_E_* < 1 + *w_I_* (Eq. 3).

### Perturbation of entire inhibitory population

For small networks consisting of a single excitatory and a single inhibitory neuron (Tsodyks et al., 1997; Litwin-Kumar et al., 2016), perturbing the inhibitory neuron will always result in a paradoxical response in an ISN. We considered whether this result holds true for larger networks with many excitatory and inhibitory neurons. We began by estimating the effect of a perturbation to the entire inhibitory population on the activity of a single inhibitory neuron (Eq. 8). We ignored any transient effect of a perturbation, comparing only the steady-state response of a network before and after the perturbation (see Materials and methods). This situation is illustrated in Fig. 1.

For the paradoxical effect to appear, a positive perturbation provided to the inhibitory population must result in a counterintuitive reduction in the activity of the inhibitory neuron under measurement. To determine whether this “paradoxical” effect occurs for a given network and given perturbation, we calculated the change in firing rate of a chosen inhibitory neuron with respect to a perturbation (see Materials and Methods).

For a stable ISN as defined above (Eq. 3; see Materials and Methods), we found that a global perturbation of the inhibitory population will always evoke a paradoxical effect. This result shows that the dynamics of our large networks are comparable to previous simplified ISN models (Tsodyks et al., 1997; Litwin-Kumar et al., 2016).

### Perturbation of a single inhibitory neuron

Since not all inhibitory neurons within a cortical region will be perturbed with electophysiological or optogen-etic approaches under realistic experimental conditions, we investigated how networks respond when only a fraction of the inhibitory neurons are perturbed. Starting with the extreme case of perturbing a single inhibitory neuron (Eq. 9), we found that a narrow range of excitatory synaptic strength *w_E_* exists, within which the paradoxical effect can be evoked (see Materials and Methods). However, the range for *w_E_* that satisfies this constraint shrinks rapidly to zero as the size of the network increases, making this regime unlikely to exist in cortex.

### Perturbation of a subset *p* of the inhibitory population

We then investigated the effect of perturbing a larger subset of the inhibitory population, as is likely to be the case under experimental conditions. We injected a positive or negative current into *p* inhibitory neurons (see Materials and Methods; Eq. 10). We found that for networks in a stable ISN regime, the relative total synaptic strength of excitatory and inhibitory neurons determines a minimum proportion *p/N* > *–λ_1_/w_I_* of the inhibitory network that must be perturbed, in order to observe a paradoxical response in the perturbed neurons (Fig. 3a). Importantly, this proportion does not depend on the size of the network *N*.

Depending on the operating regime of the network, the proportion of inhibitory neurons that must be perturbed can be considerable, approaching 100%. If a smaller proportion of the inhibitory network is stimulated, then the paradoxical response does not occur in either the perturbed or non-perturbed inhibitory neurons (Fig. 2).

**Figure 2:**
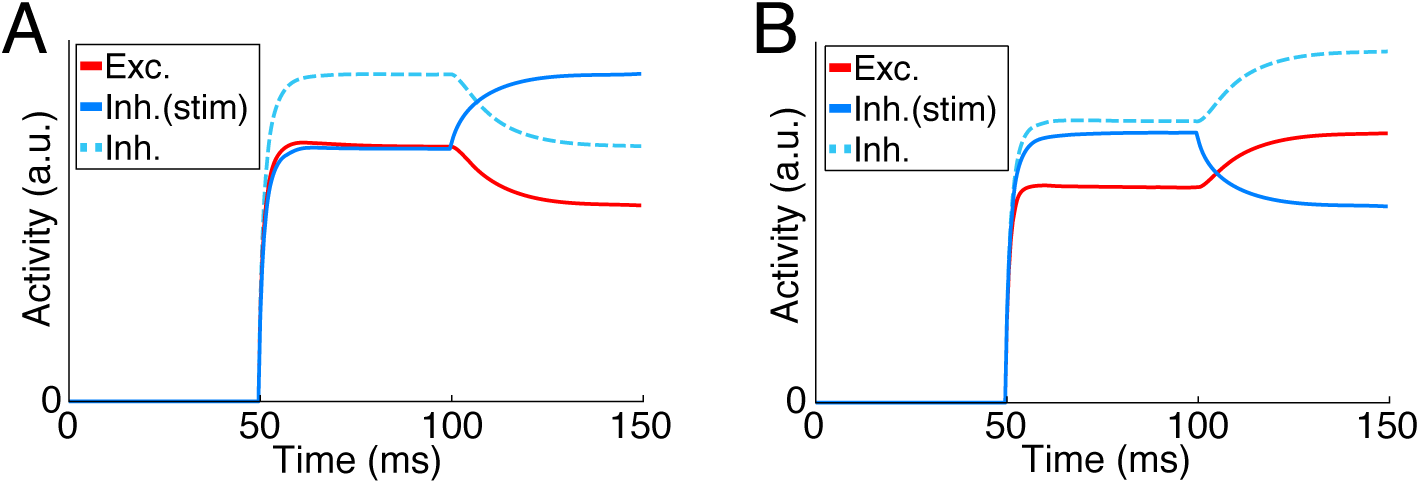
Perturbing only a proportion of the inhibitory population may not give rise to a paradoxical inhibitory response. Result of increasing (A) and decreasing (B) input to a portion *p* = 50% of the inhibitory population (compare with Fig. 1). Although this network is an ISN with same parameters as in Fig. 1, the response of inhibitory neurons to perturbation is starkly different. No particular evidence for the paradoxical response is visible. Dashed trace: response of non-stimulated inhibitory neurons, shifted up for visibility. The response of excitatory neurons (red) and non-stimulated inhibitory neurons (dashed) are identical.

**Figure 3:**
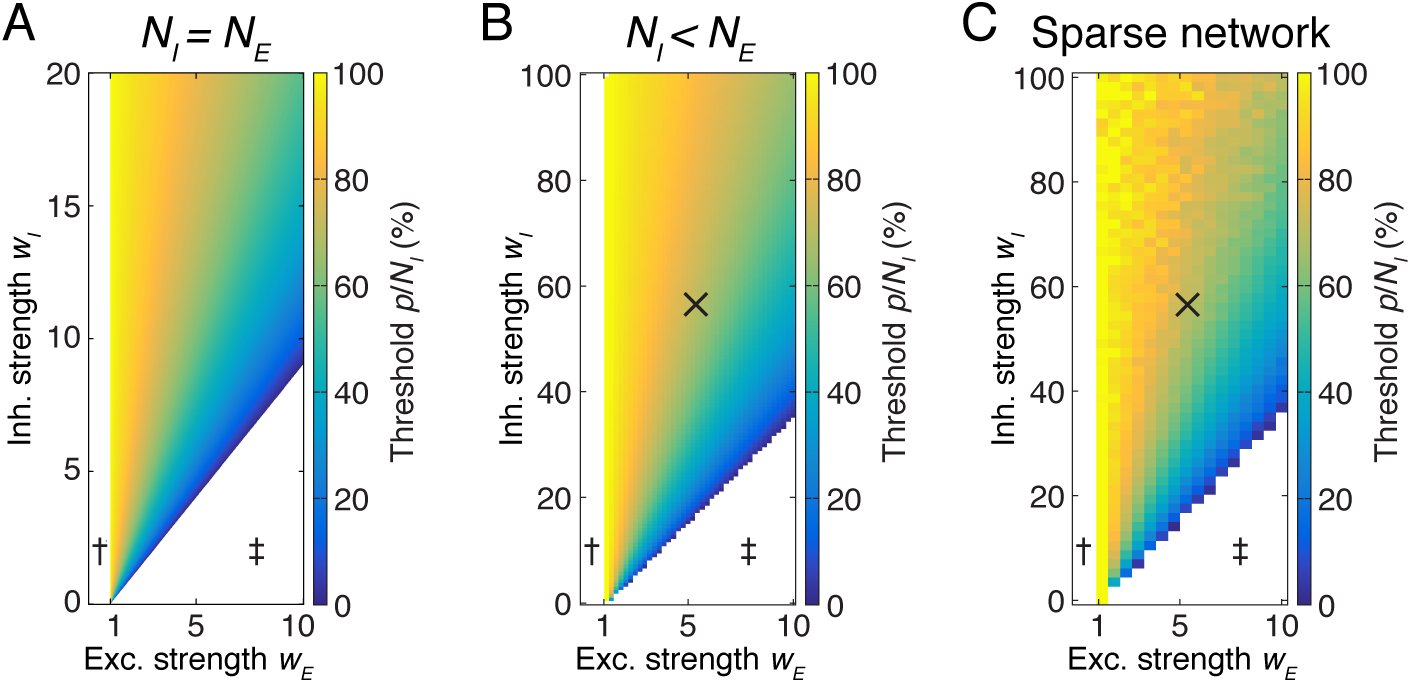
Many inhibitory neurons must be perturbed to evoke a paradoxical inhibitory response. (A) The minimum proportion of the inhibitory population *p/N* that must be perturbed under in order for the paradoxical effect to appear in the perturbed neurons, in a network with equal numbers of excitatory and inhibitory neurons. This analytical result does not depend on the size of the network *N*. Parameters: {*h_I_, h_E_, f_I_, τ*} = {1, 1, 0.5, 10 ms}. (B) The miminum proportion of inhibition *p/N_i_* for a network with *f_I_* = 20 %. Other parameters: {*h_I_, h_E_, τ, N*_E_, *N*_I_} = {1, 1, 10 ms, 80, 20}. Note the difference in scale compared with (A). (B) The minimum proportion of the inhibitory population *p/N* that must be perturbed under in order for the paradoxical effect for networks with sparse synaptic connectivity between excitatory and inhibitory neurons. Note this does not affect the overall trend for averaged response of stimulated inhibitory neurons (cf. B), but the stochastic effect of introducing sparse connections in smaller networks is evident. Parameters: {*h_EE_, h_EI_, h_IE_, h_II_*,*N_E_, N_I_*} = {0.1, 0.5, 0.5, 0.5, 4000, 1000}. × in b,c: estimated nominal parameters for mouse visual cortex {*w_E_, w_I_}* = {5.4, 56}. This estimate gives p/*N_I_* = 71 %. †: non-ISN regime; ‡: unstable regime.

### Perturbation by injecting a global inhibitory current

Some experimental perturbations, for example infusion of neurotransmitters or chemical agonists of inhibition, result in injection of inhibitory currents across the entire network (i.e. in both inhibitory and excitatory neurons). We therefore examined the case of such a global perturbation in our models (see Materials and Methods; Eq. 13). We found that this mode of perturbation cannot elicit a paradoxical inhibitory response in a network operating in a stable ISN regime. Experimental methods that globally modulate inhibitory inputs to all neurons — as opposed to perturbing the inhibitory population alone — cannot therefore be used to demonstrate an ISN regime in cortex.

### Perturbation by modifying effective inhibitory synaptic strength

It is possible that some experimental perturbations, for example infusion of a GABA antagonist, may result in a divisive rather than subtractive effect on inhibitory input. We investigated the effect of divisive perturbations by scaling the effective inhibitory synaptic strength *w_I_*. We computed the change in neuronal responses when effective inhibitory synaptic strength is perturbed, requiring that for an increase in inhibitory synaptic strength, the analogous “paradoxical” response would be for the inhibitory network to increase its activity (see Materials and Methods; Eq. 14). We provided a constant but different input current to excitatory and inhibitory neurons, 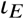 and 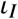 respectively.

We found that for a network operating in a stable ISN regime, there is no combination of relative excitatory and inhibitory input or synaptic weight that can give rise to a paradoxical inhibitory response when the inhibitory synaptic strength is perturbed. This result implies that global modulation of inhibitory weights, or other similar divisive modulation of inhibition, cannot be used to demonstrate an ISN regime in cortex.

### Networks with realistic proportions of excitatory and inhibitory neurons

The networks described above have equal numbers of excitatory and inhibitory neurons, similar to classical ISN networks. However, in mammalian cortex, approximately 20 % of neurons are inhibitory (Gabott and Somogyi, 1986). We therefore redefined our network following Muir and Mrsic-Flogel (2015), and set the proportion of inhibitory neurons in the network to 20 % while maintaining all-to-all non-specific connectivity. We numerically computed the proportion of the inhibitory population that must be stimulated to observe the paradoxical effect in the stimulated neurons (Fig. 3b; see Materials and Methods). In general, networks with fewer inhibitory neurons are less stable. Indeed, an increase in *w_I_* is required for stability (compare Fig. 3a with b). However, we observed the same trends for evoking a paradoxical inhibitory response in networks with fewer inhibitory neurons, as for the networks with equal numbers of excitatory input.

### Perturbations in networks with sparse connectivity

Synaptic connections between neurons in the neocortex are not all-to-all; neurons connect to their immediate neighbours with an average probability of only around 20% for recurrent excitatory connections (Gabott and Somogyi, 1986). Connections between neighbouring inhibitory and excitatory neurons are much more dense, with close to 100% connection probability between neighbouring excitatory and parvalbumin-positive inhibitory neurons (Hofer et al., 2011; Bock et al., 2011; Fino and Yuste, 2011; Martin, 2011; Bopp et al., 2014), but connection probabilities fall off dramatically with distance (Boucsein et al., 2011; see Materials and Methods).

To examine the effect of sparse connectivity we expanded upon the work in Muir and Mrsic-Flogel (2015) by introducing connection sparsity parameters that describe the number of synaptic connections made between nearby neurons, as a proportion of all possible partners. We estimated separate sparsity parameters for recurrent excitatory, exc. → inh., inh. → exc. and recurrent inhibitory connections, based on the assumption of stochastic connections formed between neurons with overlapping axonal and dendritic arbors, and to match reported connection probabilities (Peters’ rule; see Materials and Methods; Peters, 1979; Reimann et al., 2015).

By computing the proportion *p/N_I_* of the inhibitory population that must be stimulated in order to observe the paradoxical effect, we found that if one records the *average* response of stimulated inhibitory neurons then *p/N_I_* only differs from the fully-connected network in terms of stochasticity induced by the random sparsity structure of individual instances of *W* (Fig. 3c). Estimates for nominal parameters of total synaptic strength in rodent cortex are indicated by × in Fig. 3b–c, suggesting that in the order of 71 % of inhibitory contribution must be perturbed in order to observe the paradoxical inhibitory response in cortex. However, due to the spatial dependence of connectivity and the tendency for local inhibition to be strong, dense and class-specific (Bock et al., 2011; Fino and Yuste, 2011; Hofer et al., 2011; Martin, 2011; Bopp et al.,2014), inhibition may be even stronger than this estimate which is based on uniform connection probabilities. Our results predict that a large fraction of of inhibitory neurons must be perturbed to evoke a paradoxical response in cortex.

### Measuring inhibitory input currents in excitatory neurons

Litwin-Kumar et al. proposed that recording the inhibitory current received by excitatory neurons as an experimentally-accessible metric for observing the paradoxical effect of an ISN (Litwin-Kumar et al., 2016). Due to dense connectivity from the inhibitory population onto excitatory neurons (Fino and Yuste, 2011), recording net inhibitory currents provides an estimate of the mean activity of the local inhibitory population, rather than sampling from an individual inhibitory neuron. Optogenetic perturbation of the inhibitory population, while recording from individual excitatory neurons, was performed by Atallah et al. (2012). However, the behaviour of ISNs under simulated optogenetic perturbations is not known, leaving in question whether the averaging is sufficient in sparse networks, and under what conditions a paradoxical effect should be visible.

We therefore performed simulated optogenetic perturbations of the inhibitory population by injecting positive and negative currents, and recording the resulting change in inhibitory input to excitatory neurons (Fig. 4). We simulated the presence of a stimulus in the network by providing random fixed input currents to each neuron. This placed the network in a realistic regime where symmetry is broken by an input stimulus, and competition between neurons can be expressed. We then perturbed a randomly chosen proportion *p/N_I_* of the inhibitory network by providing a common input current with amplitude δ ranging (–1,1), designed to simulate perturbation by optogenetic activation or suppression.

**Figure 4:**
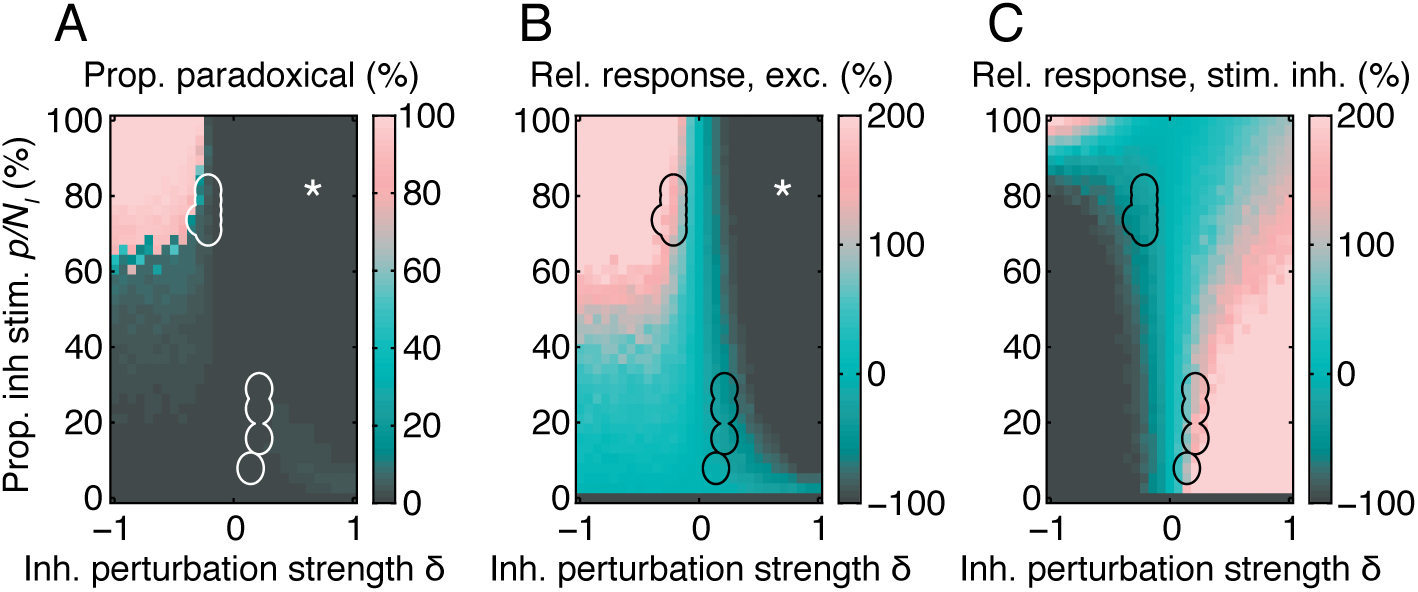
Paradoxical effects under optogenetic perturbation. (A) Responses to perturbation in an ISN-regime network, indicating the proportion of excitatory neurons that exhibit a paradoxical effect in the net inhibitory input currents, as a function of perturbation strength δ and proportion *p* of inhibitory neurons perturbed. (B–C) Relative change in (B) excitatory and (C) stimulated inhibitory neuron activity, for the same simulations as in (A). We considered that a paradoxical effect was visible when the input currents changed by at least 10% in the appropriate direction. Outlined regions in A–C indicate responses to perturbation where changes in excitatory and inhibitory activity are approximately equal to those reported by Atallah et al. (Atallah et al., 2012; see Materials and Methods). * Region where all excitatory neurons are below threshold, leading to failure of excitatory-driven inhibition. Parameters: {*w_E_, w_I_, h_EE_, h_EI_, h_IE_, h_I I_ N*_E_, *N*_I_} = {4, 100, 6.4 × 10^−3^, 0.21, 0.24, 0.99, 4800, 1200}

We recorded the amplitude of inhibitory input currents impinging on each excitatory neuron and defined an excitatory neuron as showing a paradoxical effect if inhibitory input currents were modified by at least 10% in response to the inhibitory perturbation. As shown in Fig. 4a, paradoxical effects were only observed in a substantial proportion of excitatory neurons when the majority of inhibitory neurons was inhibited. Indeed, regimes exist for ISN networks with strong excitatory and inhibitory feedback, where the paradoxical effect cannot be observed in the majority of excitatory neurons. Indicated regions in Fig. 4 correspond to the effect sizes reported in Atallah et al. (2012), as determined by comparing the relative change in firing rates of excitatory and inhibitory neurons following a perturbation (Fig. 4b, c). Under a range of choices for strengths of excitation and inhibition, the simulated perturbations equivalent in size to those reported in Atallah et al. (2012) are not sufficient to demonstrate the paradoxical effect.

### Perturbations in more realistic networks of spiking neurons

Our results so far were obtained in network models with simplified firing-rate dynamics. However, networks composed of nonlinear spiking units are known to show rich and complex activity dynamics (Brunel, 2000; Ostojic, 2014), with response properties depending on the operating regime of activity (Destexhe et al., 2003; Kuhn et al., 2004; Kumar et al., 2008). In order to verify that our results hold in more biologically realistic networks, we investigated the dynamics of paradoxical inhibitory response in networks of nonlinear conductance-based spiking neurons (see Materials and Methods).

The spiking activity of a sample network of conductance-based exponential integrate-and-fire neurons is shown in Fig. 5a, before and after perturbation of two different fractions of the inhibitory population. The perturbation was performed by *decreasing* input to a subset of inhibitory neurons.

**Figure 5:**
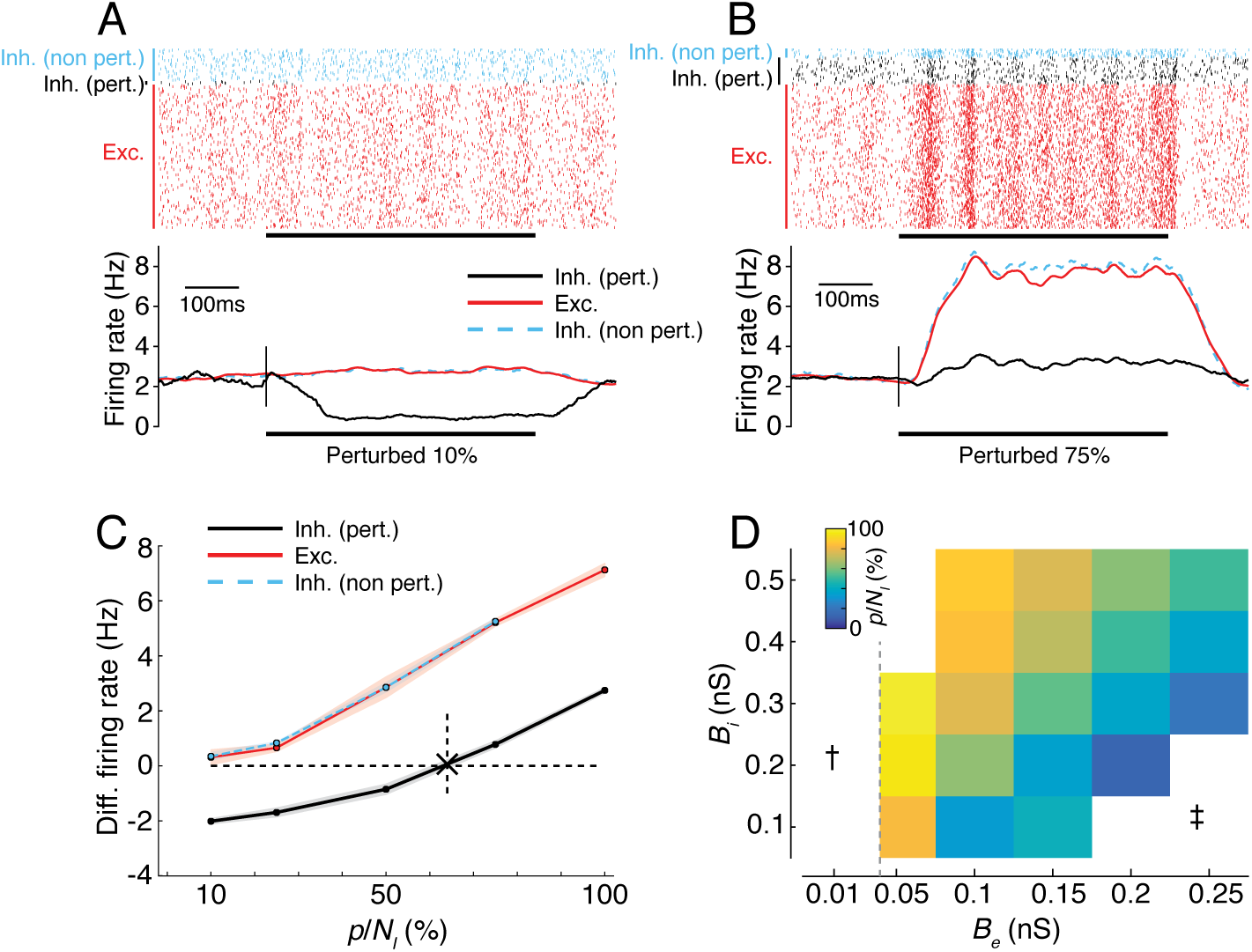
The paradoxical effect in spiking ISNs depends on the proportion of perturbed inhibitory neurons. (A–B) Result of perturbing 10% (A) and 75% (B) of the inhibitory population in a spiking network model, by reducing input to inhibitory neurons. Top: Single-trial spike rasters from the entire population. Bottom: Averaged firing rates over 10 trials (smoothed by a boxcar filter of 100 ms width). Black bar: perturbation period. c.f. Fig. 2. Red: Excitatory (Exc.) neurons; Black: Perturbed inhibitory neurons (Inh. pert.); Cyan: Non-perturbed inhibitory neurons (Inh. non-pert.). Parameters for this network: {*N_E_, N_I_, B_e_, B_i_*} = {1600, 400, 0.1 nS, 0.2 nS}. For other parameters, see Methods and Table 3. (C) Mean (dots) and std. deviation (shading) of the differential rates under a range of perturbed proportions for the network shown in A,B. Cross and dashed line in C: inferred minimum fraction of perturbed inhibition *p/N_I_* required to obtain the paradoxical effect (see Materials and Methods for details). (D) The minimum fraction *p/N_I_* for spiking networks while varying *B_e_* and *B_i_* (c.f. Fig. 3). Dashed line in D: border of the ISN regime according to a simplified linear analysis of the network (see Materials and Methods). †: non-ISN regime; ‡: unstable regime (firing rates > 100 Hz).

The average activity within each subpopulation (excitatory, perturbed inhibitory and unperturbed inhibitory) is shown in Fig. 5b. When 10% of the inhibitory population was perturbed, no paradoxical effect was observed: the (negatively) perturbed inhibitory subpopulation decreased its activity, whereas the unperturbed inhibitory and excitatory subpopulations increased their activity. However, when larger fractions (75%) of inhibitory neurons were perturbed, the network displayed the paradoxical effect by increasing the average activity of the perturbed neurons, in spite of a decrease in the input to the inhibitory network, consistent with the predictions of our firing-rate model (c.f. Fig. 2a).

To quantify the strength and presence of the paradoxical effect, we measured the average differential firing rate (perturbed rate minus baseline rate) while varying the fraction of perturbed inhibitory neurons (Fig. 5c; see Methods for details). The paradoxical effect was present when large fractions of inhibitory neurons were perturbed, as indicated by a positive differential rate. We determined the minimum fraction at which the paradoxical effect emerged by interpolating the mean differential rate, and inferring the point at which the differential rate crossed zero (Fig. 5d; see Materials and Methods). Under these simulation conditions >60% of the inhibitory neurons were required to generate a paradoxical effect.

We next examined whether the minimum fraction of inhibitory neurons *p/N_I_* required to evoke the paradoxical effect depended on the relative strengths of excitatory and inhibitory feedback, as predicted by our non-spiking simulations. To test this, we fixed all parameters of the spiking network and modified the strength of exc. and inh. conductances *B_e_* and *B_i_* (Fig. 5d). For each combination of synaptic strength, we estimated the minimum fraction of inhibition *p/N_I_* from the differential rate curves (analogous to Fig. 5c).

When excitation was too weak (white regions on the left in Fig. 5d), no paradoxical effect was visible. For these values of excitation, the network was not operating in an ISN regime, since the excitatory network alone was intrinsically stable (exc. conductance *B_e_* at and below grey vertical line; obtained from the stability analysis of the linearized network; see Materials and Methods for details). For very strong values of excitatory coupling, without sufficient inhibitory feedback (high *B_e_* and low *B_i_*), networks underwent a transition from the stable regime with low firing rates and asynchronous-irregular activity to a regime with high firing rates and large pairwise correlations. This was consistent with our analysis of firing-rate networks (cf. the unstable regime of network dynamics in Fig. 3). No paradoxical inhibitory response was observed in these unstable networks.

For intermediate values of *B*_e_, we found a smooth relationship between network parameters and the minimum fraction of perturbed inhibition *p/N_I_* required to see the paradoxical effect: networks with stronger excitation and weaker inhibition required smaller perturbations, similar to our results in firing-rate networks (cf. Fig. 5d and Fig. 3). The trend for *p/N_I_* mimicked the tendency for the network to become unstable for strong *B_e_*. The results from our spiking simulations therefore agreed well with those from our analytical and firing-rate models.

## Discussion

By examining the effects of simulated perturbations of activity in cortical network models with increasing degrees of realism, we determined what classes of perturbation could successfully detect the computational regime of cortical networks. In particular we examined the properties of *inhibition-stabilized networks* (ISNs) — networks that require inhibitory feedback in order to balance strong recurrent excitation (Tsodyks et al., 1997). This class of networks is particularly important for mammalian neocortex, since many useful computational properties — e.g. selective amplification, sharpening of tuning, noise rejection — require networks to be in an ISN regime (Douglas and Martin, 2007; Rutishauser and Douglas, 2009; Neftci et al., 2013; Muir and Cook, 2014; Hopfield, 2015).

In simple ISN models where each cell class is represented by a single unit, perturbation of the inhibitory unit reliably leads to a “paradoxical” inverse response, whereby exciting an inhibitory neuron results in a net decrease in activity (Tsodyks et al., 1997; Litwin-Kumar et al., 2016; Fig. 1). We explored whether this paradoxical response could be used to detect ISNs experimentally, by analysing larger models with many neurons, and with both homogeneous and sparse synaptic connectivity. We then tested the predictions arising from simplified firing rate models in more biologically realistic conductance-based spiking network models. We found that when inhibitory and excitatory populations are expanded, perturbing single inhibitory neurons only evokes a paradoxical response in very small networks.

In larger and more realistic networks, we found that eliciting a paradoxical inhibitory response requires a large fraction of the inhibitory population to be perturbed (Fig. 3). The proportion of cells required depends on the relative size and synaptic strengths of the excitatory and inhibitory populations, but importantly *not* on the total size of the network. For networks with parameters estimated to be similar to mouse visual cortex, we found a large majority of inhibitory neurons must be perturbed to evoke a paradoxical response (>70%; Fig. 3b). Interestingly, connection sparsity does not affect the average minimum proportion of the inhibitory network that must be perturbed (Fig. 3c). Therefore, dense inhibitory feedback and sparse excitatory recurrence as present in mammalian cortex (Hofer et al., 2011; Bock et al., 2011; Martin, 2011; Bopp et al., 2014) does not imply that an ISN regime should be a straightforward observation. Our results suggest that establishing whether cortical networks operate in the ISN regime requires application of optogenetic strategies that allow perturbation of the vast majority of inhibitory interneurons in the circuit.

### Factors underlying the paradoxical effect in network models

Simplified network models (as in Tsodyks et al., 1997 and Litwin-Kumar et al., 2016) display robust paradoxical effects in response to perturbations of the inhibitory system. Since these networks use single neurons to represent the entire inhibitory population or entire inhibitory classes, they implicitly assume that global or class-global perturbations are made to the network. Our results imply that this assumption is crucial to their results; we showed that networks operating in an ISN regime will not display a paradoxical inhibitory response unless a minimum proportion of the inhibitory population is perturbed (Fig. 3). Care is therefore needed in interpreting these earlier results in light of the complex inhibitory system in cortex.

We found that including sparsity in local recurrent connectivity did not change the minimum proportion of the inhibitory population that must be perturbed to evoke a paradoxical response (Fig. 3c). This is because the effects of sparse connectivity average out as the network size increases. Although the local minimum proportion of inhibitory neurons fluctuates across the network under sparse connectivity, we found that if the average total excitatory and inhibitory synaptic strength per neuron is held fixed, the average minimum proportion is then identical between fully- and sparsely-connected networks.

### Application to experimental methods for inhibitory perturbation

#### Electrical stimulation

The activity of a neuron can be conveniently perturbed electrically by passing positive or negative currents through a recording electrode. However, since only small numbers of cells can be perturbed simultaneously using electrophysiological methods, our results imply that paradoxical responses will not be observed in cortex even if an ISN regime exists (Fig. 3).

#### Chemical stimulation

Several agonists and antagonists of GABA receptors exist, with varying selectivity for receptor subtypes (Chebib and Johnston, 1999; Krall et al., 2015). Ant/agonists that result in additive or subtractive modulation of inhibition are equivalent to adding or removing activity from both inhibitory and excitatory neurons. Our results for network-global perturbations of input inhibitory currents imply that ant/agonists with this mechanism of action cannot induce a paradoxical inhibitory response regardless of the presence of an ISN regime (Eq. 13).

Ant/agonists that instead result in multiplicative or divisive modulation of inhibitory input currents are equivalent to a modification of inhibitory weight. Our results for network-global modifications of effective inhibitory weight showed that this type of perturbation also cannot induce a paradoxical inhibitory response (Eq. 14).

#### Optogenetic perturbation

Optogenetic approaches enable photo-activation or -suppression of specific neuron populations through genetically targeted expression of light sensitive proteins (Boyden et al., 2005; Han and Boyden, 2007; Zhang et al., 2007; Aston-Jones and Deisseroth, 2013). This approach was taken by Atallah et al. to stimulate and suppress activity in parvalbumin positive (PV+ve) inhibitory neurons, coupled with simultaneous *in vivo* electrophysiology to record responses to stimulation in individual excitatory and inhibitory neurons (Atallah et al., 2012). Atallah et al. showed that mild perturbation of PV+ve neurons (approximately-40% suppression and +20% activation; their Fig. 2) did not modify tuning of stimuli in mouse V1 (Atallah et al., 2012). The resulting changes in excitatory activity were also mild, and inhibitory currents received by excitatory neurons did not show a paradoxical effect on average (their Fig. 5).

Our findings cast new light on these results by showing that a large majority of inhibitory neurons must be perturbed to evoke a paradoxical response (Fig. 3). It is therefore not surprising that Atallah et al. did not observe such an effect, especially considering that PV+ve inhibitory neurons comprise less than 50% of inhibitory neurons in the superficial layers of cortex (Markram et al., 2004; Gonchar et al., 2007) and a similar proportion of inhibitory synapses (Binzegger et al., 2004), placing a hard upper bound on the proportion of inhibitory neurons available for perturbation in their experiments.

We also showed that measuring inhibitory currents received by excitatory neurons (Litwin-Kumar et al., 2016) does not guarantee a paradoxical effect will be observed in sparsely-connected ISNs. In Fig. 4, white outlines mark regimes of inhibitory perturbation that match the effects on excitatory and inhibitory activity observed by Atallah et al. (2012). In the presence of strong inhibition and sparse excitatory feedback in cortex, only a minority of excitatory neurons is expected to show a paradoxical effect in inhibitory input currents. The lack of a paradoxical change in inhibitory input currents observed by Atallah et al. therefore does not rule out the presence of an ISN regime in rodent cortex. Our results suggest that photosuppression of inhibitory neurons can be used to detect an ISN regime, but that optogenetic transducer proteins must be expressed in a majority of inhibitory neurons to do so. This could be achieved using an interneuron-specific promoter such as glutamate decarboxylase (GAD) to target all cells that synthesize GABA. Large area photostimulation could then be used to inhibit a large fraction of inhibitory neurons, rather than the subpopulation studied in Atallah et al̤ The presence or absence of a paradoxical effect could then be determined by examining the inhibitory drive onto pyramidal cells (Litwin-Kumar et al., 2016).

#### Other evidence for the operating regime of cortex

Surround suppression in cat visual cortex is consistent with an ISN operating regime, under the assumption that projections from the visual surround specifically modulate the inhibitory population (Ozeki et al., 2009). Robust propagation of oscillatory activity in several species (Timofeev et al., 2000; Rubino et al., 2006; Wu et al., 2008; Stroh et al., 2013) suggest that recurrent excitation is strong enough to regenerate activity (Beurle, 1956; Compte et al., 2003; Wu et al., 2008). In the rodent, supralinear amplification of single spikes (London et al., 2010) provides additional evidence for strong excitatory recurrence in cortex. More directly, anatomical and physiological estimates of synaptic contributions from various neuronal classes place both cat and rodent cortex in an ISN regime (Binzegger et al., 2004; Lefort et al., 2009; Binzegger et al., 2009).

#### Limitations of our results

We examined networks in which local excitatory connections were made sparsely, but with identical probability between all excitatory neurons. However, recurrent excitatory connectivity is biased by functional similarity in both rodent non-columnar visual cortex (Ko et al., 2011; Cossell et al., 2015) and columnar visual cortex (Malach et al., 1993; Bosking et al., 1997; Muir et al., 2011; Martin et al., 2014). We examined perturbations in networks with recurrent excitatory selectivity (Muir and Mrsic-Flogel, 2015) but found this did not affect the perturbation required to evoke a paradoxical inhibitory response. We included only a single inhibitory class in our networks, compared with the multiple classes present in cortex; however recent modelling results show that the dynamics and presence of ISN regimes are similar in networks with multiple inhibitory classes (Litwin-Kumar et al., 2016).

Our results illustrate that emergent dynamics in the highly recurrent networks of mammalian neocortex can complicate experimental detection of the network configuration. In particular, intuitions derived from small schematic models about how classes of neurons interact may not hold in more realistic networks. Our model-based predictions show that while it is possible to test for an ISN regime in cortex using optogenetics, particular experimental conditions are required to do so successfully. Computational modelling of cortical dynamics is therefore an essential tool to predict the effect perturbations will have under particular hypotheses of cortical interactions, and for guiding experimental design to test those hypotheses.

## Materials and methods

### Neuron and network dynamics

We begin by defining a simple model for a cortical network containing equal numbers of excitatory and inhibitory linear-threshold neurons (Wilson and Cowan, 1973). The activity dynamics of the network evolve according to the system of equations

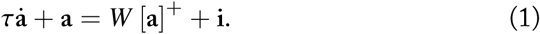

Here, *τ* is the activation time constant applied to all neurons in the network; a =(*x_1_*,*x_2_*,…,*x_N_*,*y_1_*,*y_2_*,…,*y_N_*)*^T^* is the vector of instantaneous activations (i.e. total input current in Amps) of excitatory neurons *x_i_* and inhibitory neurons *y_i_* at time 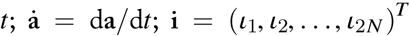is the vector of instantaneous input currents applied to each neuron; the notation [·]^+^ indicates the linear-threshold current to firing rate (I-F) transfer function [*x*]^+^ = max (*x*, 0); and *W* is the weight matrix of the network. *W* is expressed in units of ^A^/Hz, and includes any required I-F gain factors.

### Homogeneous networks with equal numbers of excitatory and inhibitory neurons

With the firing rate of each neuron evolving under the dynamics given in Eq. 1 above, we define a network weight matrix *W* with dimensions 2*N ×* 2*N*, given by

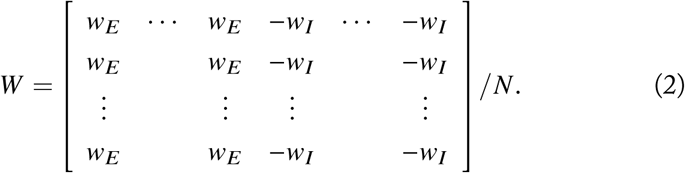

In this network, the first *N* neurons are excitatory and the subsequent *N* inhibitory, with homogenous all-to-all connectivity. More cortically-realistic network structures will be examined below. Neuron gains are assumed to be incorporated into the weight matrix *W*.

### Stability and fixed-point response analysis

We examine the fixed points and stability of the network defined in Eq. 2 evolving under the dynamics in Eq. 1, linearized in the partition where all neurons are active (Hahnloser, 1998; Muir and Cook, 2014). The stability of these networks is determined by examining the eigenvalues and trace of the system Jacobian *J* = (W – *I*)./*τ*, where *I* is the 2*N* × 2*N* identity matrix. Networks of this structure have a trivial eigenvector **1***^T^* corresponding to the eigenvalue (*w_E_* – *w_I_ – 1)/τ = λ_1_/τ*. The trace of the Jacobian is given by Tr [*J*] =*w_E_* – *w_I_* – 2*N*)/*τ* To guarantee that the network is stable under any finite input (i.e. *bounded-input/bounded-output* or BIBO stability), the eigenvalue *λ*_1_ < 0. We therefore obtain an upper bound on the total weight *w_E_* provided by each excitatory neuron relative to the strength of inhibition, given by *w_E_* < 1 + *w_I_*. The system trace provides an additional stability constraint *w_E_* < 2*N* + *w_I_*, which for these networks is always a looser bound than that imposed by λ*_I_* < 0. For the network to require inhibitory feedback for stability, the excitatory network alone must be unstable; that is, when *w_I_* = 0. This introduces a lower bound on excitatory feedback *w_E_* > 1. For a stable ISN, we therefore obtain the following constraint relating excitation and inhibition:

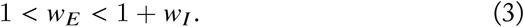

We analyse the response of the network in steady state, where a constant input is provided and the system allowed to come to rest. The fixed point response of the network is obtained by solving the system dynamics in Eq. 1 for the condition 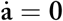 for an input i, and is denoted 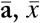 and 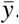. For a single neuron *j*, the fixed point is given by

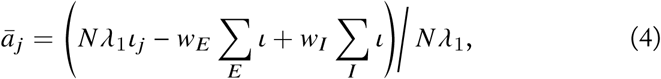
where 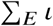 and 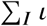 denote a summation of the input currents provided to all excitatory or inhibitory neurons respectively, and *λ*_1_ = *w_E_* – *w_I_* – 1 as defined above.

### Homogenous networks with unequal numbers of exc. and inh. neurons

We additionally define networks with varying proportions of inhibitory neurons *f_I_* (Muir and Mrsic-Flogel, 2015). In this work we examine networks where *f_I_* = 0.2, while maintaining all-to-all non-specific connectivity (i.e. in the notation of Muir and Mrsic-Flogel 2015: *h_E_, h_I_* = 1; *M* = 1; *κ* = ∞). In these networks, *N_I_* = *N f_I_* and *N_E_ = N* (1 – *f_I_*) denote the number of inhibitory and excitatory neurons respectively. Stability and fixed point response analysis are performed following the procedures above.

### Networks with sparse connectivity

To generate sparse networks we follow the procedures in Muir and Mrsic-Flogel (2015). Briefly, fully-connected network weight matrices *W* are combined with a sparse *N × N* boolean matrix *D*. To generate *D*, the appropriate number of non-zero elements for a column, as defined by *h* and *N*, are distributed randomly within each column. The network weight matrix is then given by *W*’ = *D* o W, where o denotes the elementwise Hadamard or Schur product, and *W*’ is renormalized such that columns of *W*’ sum to *w_E_* and *w_I_*. In the limit as *N* → ∞, the elements of *D* can be assumed to be independent, and therefore approximated by a Bernoulli distribution. This assumption assists in estimating the eigenvalue spectrum radius of *W*’, described below.

### Estimating the sparsity of connections in cortex

To estimate realistic parameters for the sparsity of local connections in cortex, we assume that connections between neurons are made stochastically according to the overlap of simulated axonal and dendritic densities, which are modelled as 2-dimensional Gaussian fields. The overlap between two 2-dimensional Gaussian fields is proportional to

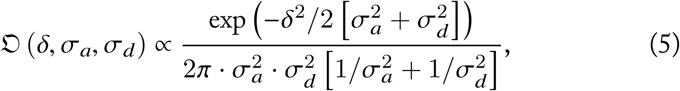
where δ is the 2-dimensional Euclidean distance between two points, and the standard deviations of axonal and dendritic fields are given by *σ_a_* and *σ_d_*: spectively. Eq. 5 is used to compute connection probability fields as a function of axonal and dendritic spreads.

We define the notation 〈·〉 to indicate that the quantity within the brackets should be normalized such that is forms a p.d.f. over 2-dimensional space; that is, 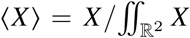. The synapse formation probability from neuron class *A* to class *B* is then given by

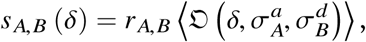
where *A* and *B* are either *E* or *I* for excitatory and inhibitory, and *r_A,B_* is the proportion of synapses from class *A* that target class B. The factors *r_A,B_* allow us to incorporate class-specific connectivity, which appears to exist in mouse visual cortex in the connections from excitatory to inhibitory neurons (Bock et al., 2011; Bopp et al., 2014).

We define the expected number of synapses from class *A* to class *B* as *n_A,B_* (δ) = *SA · s_A,B_* (δ). The connection probability *p_A,B_* from a neuron of class *A* to a neuron of class *B* at a distance δ is then given by

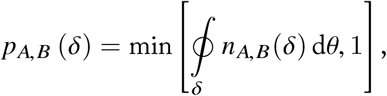
where 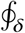d*θ* indicates integration around an annulus of distance δ from the origin (see Fig. 6). The parameters given in Table 1 result in a proximal *E → I* connection probability of *pE, I* ≈ 90%, and proximal *E → E* connection probability of *PE*,*E* ≈ 25% (see Fig. 6).

**Figure 6:**
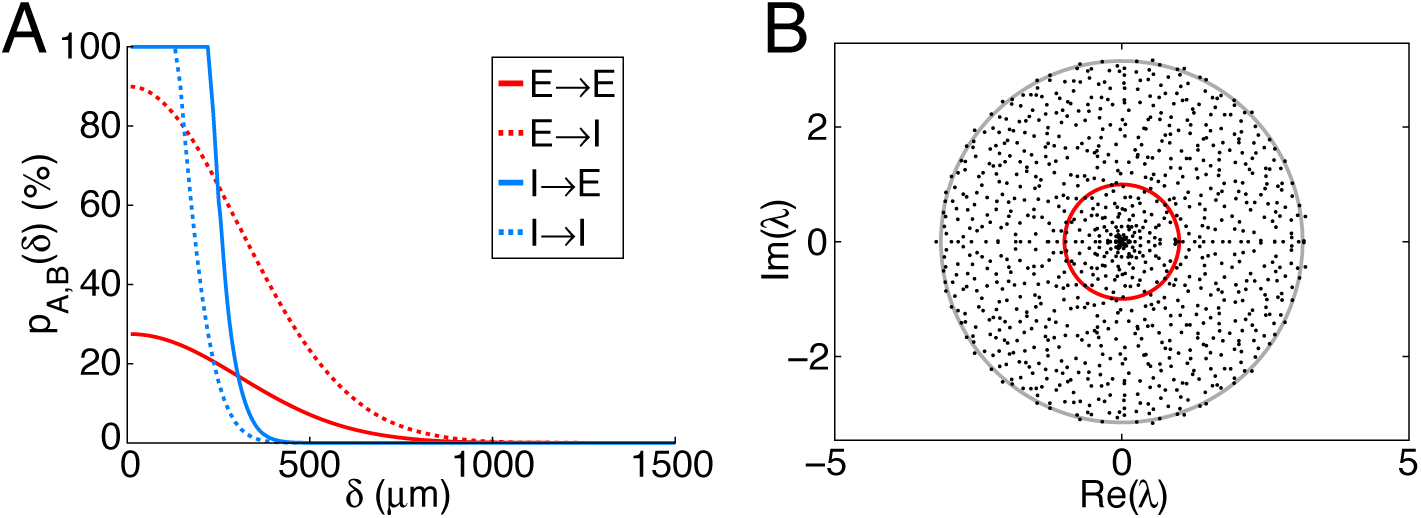
(A) Simulated connection probability *p*_A,B_ (*δ*) between neuron classes *E* and *I*. Parameters given in Table 1. (B) Eigenvalue spectrum for a sparse network with {*w_E_, w_I_, h_EE_, h_EI_, h_EI_, h_I I_, N*} = {5.4, 56, 0.022, 0.072, 0.084, 0.34, 1000}. The trivial eigenvalue at λ = −8 is not shown. Unit circle (red) and expected bulk radius *r*b (grey; Eq. 7) shown for reference.

**Table 1:** Parameters for estimating connection sparsity.

| Parameter |  | Value | Refs. |
| --- | --- | --- | --- |
| Axonal width | $4\sigma_E^a, 4\sigma_I^a$ | 1200 $\mu\text{m}$, 300 $\mu\text{m}$ | |
| Dendritic width | $4\sigma_E^d, 4\sigma_I^d$ | 300 $\mu\text{m}$, 300 $\mu\text{m}$ | |
| No. axonal syn. in L2/3 | $S_E, S_I$ | 8142, 8566 | Binzegger et al., 2004* |
| Density of neur. in L2/3 | $\eta$ | 36000 $\text{mm}^{-1}$ | Schüz and Palm, 1989 |
| Prop. neur. class A | $f_E, f_I$ | 80 %, 20 % | Gabott and Somogyi, 1986 |
| Prop. of $A \rightarrow B$ syn. | $r_{E,I}$ | 45 % | Bock et al., 2011; Bopp et al., 2014 |
| | $r_{E,E}$ | $1 - r_{E,I}$ | † |
| | $r_{I,E}$ | $1 - f_I$ | † |
| | $r_{I,I}$ | $f_I$ | † |

Abbreviations: neur. neurons, syn. synapses; prop. proportion; $E$ exc. excitation, excitatory; $I$ inh. inhibition, inhibitory. \* Including an estimate of double the number of synapses per neuron in mouse cortex compared with cat cortex. † i.e. Non class-specific connectivity.

The sparsity (and equivalently, the fill factor *h*) of connections from class *A* to class *B* is therefore estimated by

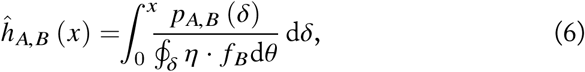
where Eq. 6 should be integrated out to a distance *x* at which the connection probability drops to zero. Taking *x* = 1500 *μ*m for excitatory neurons and *x* = *750 μ*m for inhibitory neurons, we estimate 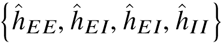 = {0.022, 0.072, 0.084, 0.34}. These low fill factors make the resulting network instances highly unstable, even in the presence of strong inhibitory feedback, due to expansion of the eigenspectrum bulk (Muir and Mrsic-Flogel, 2015; see Fig. 6). The expected radius *qb* of the eigenspectrum bulk for a network with class-dependent fill factors is given by

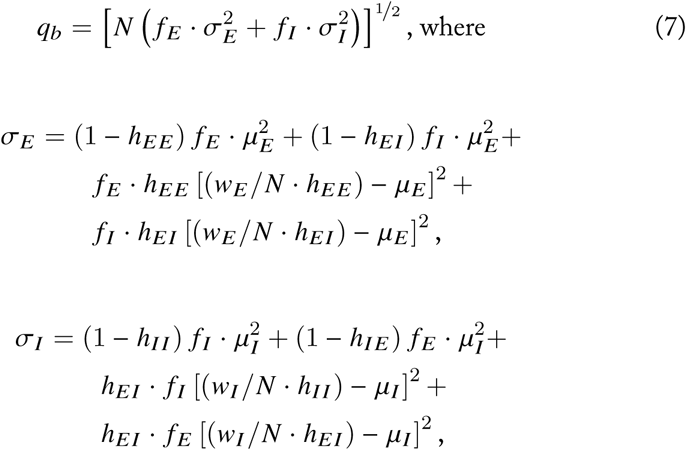

*μ_Ε_* = *W_E_/N* and *μ_I_* = *w_I_/N* (c.f. Muir and Mrsic-Flogel, 2015). To ensure stability in networks with scale smaller than cortex itself, we therefore simulate networks where the radius of the bulk eigenspectrum is controlled by scaling *h_*_* by a common factor, such that *qb* ≈ 1.

## Perturbation framework

In general, we introduce a perturbation to a network by defining an input *s* (δ), where *s* defines the input currents to all neurons in a network and δ is a small perturbing effect (δ > 0 corresponds to a positive perturbation in input and δ < 0 corresponds to a negative perturbation. For example,

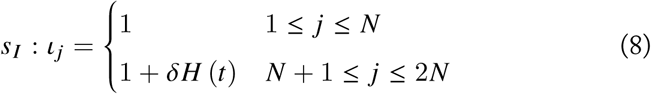
defines a scheme where all neurons receive a constant input (“1”), and the entire inhibitory population (*N* + 1 ≤ *j* ≤ *2N*) receives an extra perturbing input δ at *t* = 0. Here, *H* (*t*) is the Heaviside step function.

We assume that a perturbation is made in a network where every neuron is active; inactive subsets of the network can be removed entirely from the system (Hahnloser, 1998; Muir and Cook, 2014). We examine the fixed point 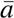 (Eq. 4) of the analytical network, linearized in the state partition when all neurons are active (Muir and Cook, 2014). We assume that the perturbation δ is small enough that no neuron is pushed below threshold.

We assume that a perturbation is only made once the transient response of the network has settled and the network has reached a stable fixed point. We therefore examine the mean-field fixed-point response of these networks, under the assumption that the effect of stochastic or oscillatory dynamics will be removed by averaging. We likewise neglect the transient effect of a perturbation, and examine only the resulting fixed point response subsequent to the perturbation (i.e. at *t* =∞).

Following a perturbation, we examine the difference between perturbed and unperturbed inhibitory activity *s* : 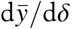, under a given perturbation s. Generally, we look for a “paradoxical” response of inhibition such that *s* : 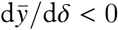 for δ > 0. For example, under the perturbation of the entire inhibitory population defined in Eq. 8 above, the change in inhibitory activity in response to the perturbation is given by

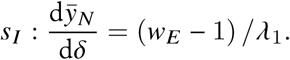

For this response to the perturbation to meet the characteristics of a paradoxical inhibitory response, we require that 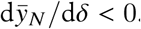. Combining this requirement with the conditions for a stable ISN (Eq. 3), we obtain the constraints on network configuration that ensure a paradoxical inhibitory response is observed in a stable ISN. By doing so, we find that the constraints already required by Eq. 3 guarantee that a paradoxical inhibitory response will be observed under the global inhibitory perturbation *s_I_*. This result implies that a stable ISN will always display a paradoxical response when the entire inhibitory population is perturbed.

### Perturbation of a single inhibitory neuron

We examined the other extreme of perturbing a single inhibitory neuron, such that

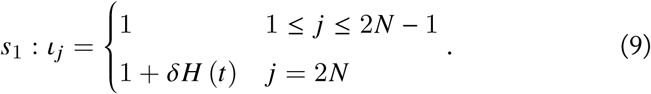

As before, we computed the change in fixed-point response of a single inhibitory neuron, when that neuron is perturbed, given by

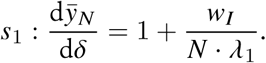

Under the requirement that a perturbation must lead to a paradoxical response (i.e. *s*_1_ : 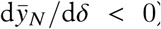), we find an additional constraint on the excitatory weight *w_E_* > 1 + *w_I_* (*N* − 1) */N*. This implies that a stable ISN can exhibit a paradoxical effect when a single inhibitory neuron is perturbed, iff 1 + *w_I_* (*N* − 1) /N < *w_E_* < *1* + *w_I_*. We note that (*N* − 1) */N* → 1 as *N* → ∞, and therefore the range for *w_E_* that satisfies this constraint approaches zero with increasing *N*.

### Perturbation of a subset *p* of the inhibitory population

We investigated the effect of perturbing a subset *p* of the inhibitory population, defined by

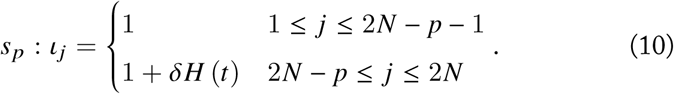

The derivative of fixed point activity is then given for perturbed inhibitory neurons by

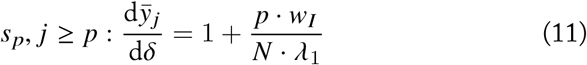
and for non-perturbed inhibitory and for excitatory neurons by

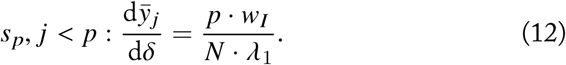

Under the constraint *s_p_, j* ≥ *p* : 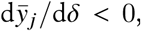, Eq. 11 implies that at least a proportion *p/N* > −*λ_1_/w_I_* of the inhibitory population must be perturbed in order to observe a paradoxical effect in the perturbed neurons.

### Perturbation by injecting a global inhibitory current

We examined the effect of perturbing the entire network by injecting a global inhibitory current, as might be produce by infusing cortex with a GABA agonist. The perturbation is defined by *S_g_* : 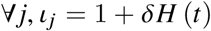. The derivative of fixed point activity for all neurons is then given by

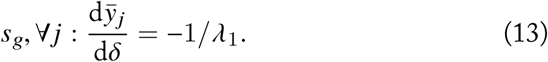

Since *S_g_* : 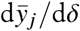 is always positive for a stable ISN (i.e. satisfying Eq. 3), no paradoxical response of inhibitory neurons is possible under the network-global perturbation *S_g_*.

### Perturbation by modifying inhibitory weight *w_I_*

Alternatively, infusion of GABA agonists or antagonists might result in an divisive rather than subtractive effect on inhibitory input currents. We therefore computed the change in fixed point response 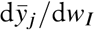 when the total inhibitory weight *w_I_* is perturbed, requiring that for an increase in inhibitory weight, the paradoxical response would be for the inhibitory network to increase its activity: i.e. 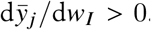. We define the input to the network

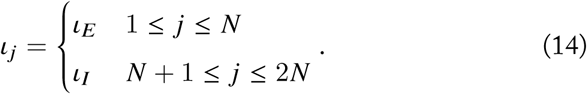

The fixed point response of the network under this input is given by

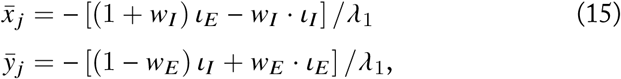
and the resulting change in fixed point response by

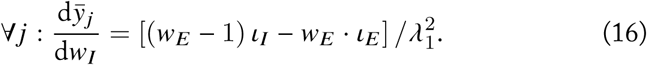

For a stable ISN, a regime exists such that if the inputs to excitatory and inhibitory neurons differ 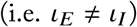, then the paradoxical response 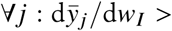 is evoked when 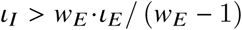. Unfortunately this regime only occurs when 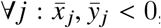, that is when the network is silenced.

### Comparison with optogenetic perturbation results from Atallah et al. (2012)

Atallah et al. used optogenetic activators and inhibitors, expressed selectively in parvalbumin-positive inhibitory neurons, to perturb inhibitory activity in mouse visual cortex (Atallah et al., 2012). They recorded responses to visual stimuli of varying contrast in the presence of optogenetically induced inhibitory suppression and activation, while recording inhibitory and excitatory synaptic input currents impinging on excitatory neurons. For comparison with these results, we found combinations of network and perturbation parameters in our simulated networks that resulted in similar perturbations of inhibitory and excitatory activity observed by Atallah et al. (Fig. 4).

We defined a simulated excitatory neuron as displaying a paradoxical response if the result of an inhibitory perturbation was to shift the net inhibitory input current by at least 10% of its unperturbed value.

## Spiking networks with conductance-based neurons

### Neuron model

Spiking neurons were modelled using an exponential integrate-and-fire model (Brette and Gerstner, 2005), without adaptation. The dynamics of the membrane potential *V_m_* (*t*) of a single model neuron evolved under the equation

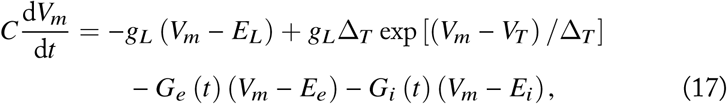
where *C* is the membrane capacitance, *g_L_* is the leak conductance and *E_L_* is the resting potential. The exponential term describes the activation of sodium current. The parameter Δ*_T_* is called the slope factor and *V_T_* is the threshold potential. Once the membrane potential *V_m_* reaches the threshold *V_T_*, a spike is emitted and the membrane potential is reset to a fixed voltage, *V*_reset_, for a refractory period *t*_ref_.

*E_e_* and *E_i_* are the reversal potentials for excitation and inhibition, respectively. *G_e_* (*t*) and *G_i_* (*t*) represent the total excitatory and inhibitory conductances at time *t*, given by

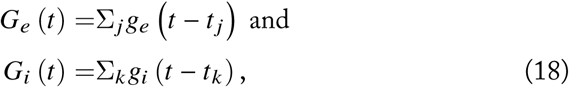
where the times of occurrence of excitatory and inhibitory synaptic events are denoted by *t_j_* and *t_k_*, respectively. *g_e_* and *g_i_* denote the membrane conductance changes elicited by a single excitatory or inhibitory synaptic event, which are modelled as alpha-functions, given by

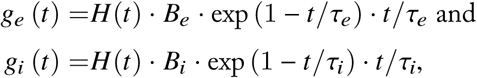
where *B_e_* and *B_i_* denote the peak excitatory and inhibitory synaptic conductances, respectively. The integral of the conductances is given by

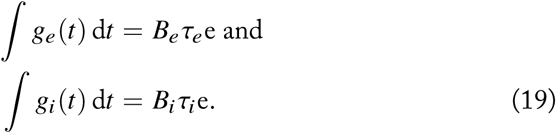

The default parameters of the neuron model are listed in Table 2. Default values of peak synaptic conductances (as in Fig. 5a-c) were *B_e_* = 0.1 nS, *B_i_* = 0.2 nS, and *τ_e_* = 1 ms, *τ*_i_ = 1 ms. Note that the effective time constant of the synapses, defined as the time from a spike until the synaptic current decays to the 10% of the peak current, is much longer (*τ*_eff_ = 4.9 ms for *τ* =1 ms). To simulate the spiking networks, we used the NEST software (Gewaltig and Diesmann, 2007). The implementation uses a fourth order Runge-Kutta-Fehlberg solver with adaptive step size to integrate the differential equation.

**Table 2:**
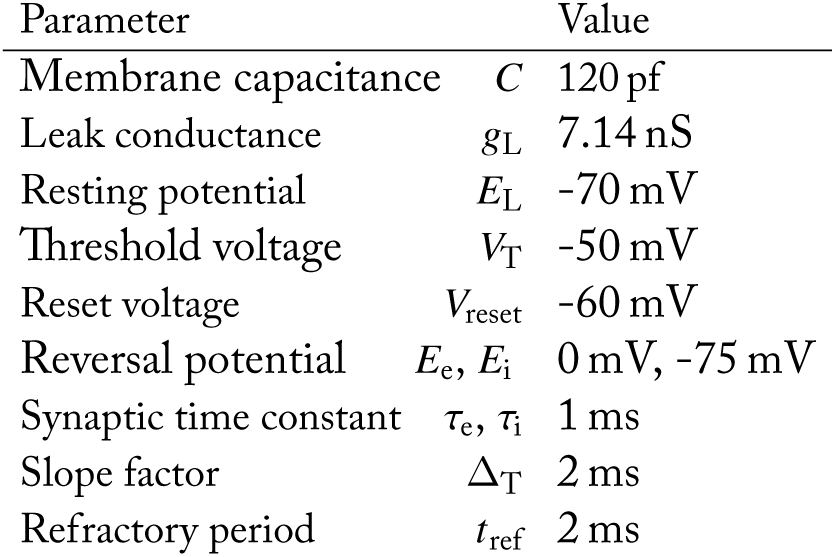
Parameters of the spiking neuron model.

### Network simulations

Networks were composed of *N_E_* excitatory and *N_I_* inhibitory neurons. Excitatory and inhibitory neurons had the same properties and parameters as described above. All neurons received a baseline input. This was modelled as an independent homogeneous Poisson process with firing rate *r_b_*. The strength of synaptic connectivity is parameterized by the peak synaptic conductance, which was denoted as *B_b_* for the baseline input. Connection delays were chosen as the fixed value of *d* for the input synapses; synaptic delays for recurrent connections were drawn from a random exponential distribution with mean *d*.

Recurrent connections were drawn from a binomial distribution. The mean connection probability from the pre-synaptic subpopulation *X* ∈ {*E, I*} to post-synaptic subpopulation *Y* ∈ {*E, I*} was *C_x→y_*. The connection weights between established connections were drawn from a truncated Gaussian distribution with a mean of *B_x→y_* and standard deviation of *B_x→y_*/5. The mean value for E → E and E → I connections were set to *B_E→E_* = *B_E→I_* = *B_e_;* similarly, the mean value for I → E and I → I connections were set as *B_I→E_* = *B_I→I_* = *B_i_*. The parameter space for the balance of excitation and inhibition in the network is scanned by changing these two parameters (e.g. in Fig. 5d).

The stimulation protocol of the network comprized of three phases: an initial transient phase where the spiking activity was not analysed (*T*_trans_); the baseline duration where the normal activity of the network was recorded (*T*_base_); and the perturbation period during which a certain fraction of the inhibitory population was perturbed (*T*_pert_). To obtain reliable estimates of firing rates, simulated perturbations were repeated for *N*_trial_ trials, with each trial lasting for *T*_trial_ = *T*_trans_ + *T*_normal_ + *T*_pert_. The default parameters of network simulations are listed in Table 3.

**Table 3:** Parameters of spiking network simulations

| Parameter |  | Value |
| --- | --- | --- |
| Number of neurons | $N_E, N_I$ | 1600, 400 |
| Connection probability | $C_{E \rightarrow E}, C_{E \rightarrow I}, C_{I \rightarrow E}, C_{I \rightarrow I}$ | 15 %, 15 %, 100 %, 100 % |
| Baseline input | $r_b$ | 9.6 kHz |
| Strength of baseline input | $B_b$ | 0.1 nS |
| Average synaptic delay | $d$ | 0.1 ms |
| Simulation time |  |  |
| (transient, baseline, perturbation) | $T_{\text{trans}}, T_{\text{base}}, T_{\text{pert}}$ | 0.15 s, 0.5 s, 0.5 s |
| Number of trials | $N_{\text{tr}}$ | 5 (Fig. 5d) or 10 (Fig. 5a-c) |
| Size of input perturbation | $\delta$ | 0.4 kHz |
| Size of perturbed inhibitory subpopulation | $p$ | {40, 100, 200, 300, 400} |

The perturbation was performed by reducing the baseline input to *p* inhibitory neurons by δ = 0.4 kHz (i.e. by ∼ 4%), and it was repeated for a range of inhibitory fractions *p/N_I_* = {0.1, 0.25,0.5,0.75,1}. For each perturbation, the mean firing rates of each subpopulation (excitatory, non-perturbed inhibitory and perturbed inhibitory) in the normal state (*r*_base_) and during perturbation (*r*_pert_) were computed by averaging over time, trials and the subpopulation. The change in the firing rate due to perturbation was then computed as rdiff = *r*_pert_ – *r*_base_. As the perturbation is performed by *decreasing* the input to a fraction of inhibitory subpopulation, a positive *r*_diff_ for the perturbed inhibitory fraction implies the existence of the paradoxical inhibitory response. We estimated the minimum fraction of inhibition to see this paradoxical effect for a given network (i.e. the value of *p/N_I_* such that *r*_diff_ = 0) by linearly interpolating *r*_diff_.

### Mean-field approximation

The mean-field analysis of the network dynamics was performed by analysing the average behaviour of the network. Let *r_e_* and *r_i_* denote the mean rates of the excitatory and inhibitory populations within a network. Combining Eq. 18 and Eq. 19, the temporally averaged excitatory and inhibitory conductances input to an example neuron can be written as

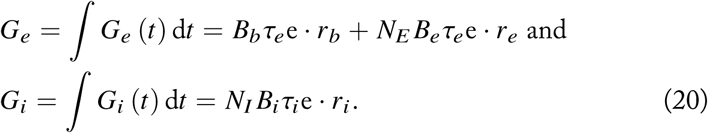

The total excitatory conductance *G_e_* is composed of two terms: the baseline external input, and recurrent input from presynaptic excitatory neurons. The inhibitory conductance *G_i_* results from presynaptic inhibitory neurons in the network.

To obtain the effective change in the membrane potential as a result of these input conductances, we must consider the effective drives from Eq. 17. We write

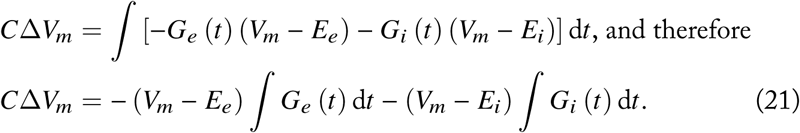

Here, we have made a simplifying assumption that the population-average membrane potential of the network is constant, and can be approximated by the time-averaged membrane potential of the network, denoted by *V_m_*. Substituting Eq. 20 into Eq. 21 we obtain the effective change in membrane potential *V*_tot_, given by

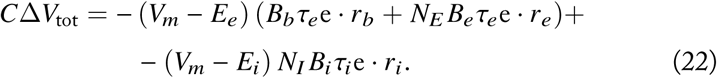

Note that the effective input is similar for any neuron, independent of its subtype identity (excitatory or inhibitory). Furthermore, we make the *ansatz* that the rates of excitatory and inhibitory subpopulations are the same: *r_e_* = *r_i_* = *r*. This is based on the fact that both subtypes have the same single cell parameters and network connectivity profiles, and the input to both subnetworks is similar in the unperturbed state. Due to this homogeneity, they have the same mean firing rates. Eq. 22 can therefore be further simplified to

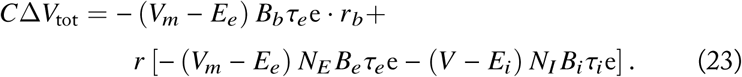

The first term on the right hand side is a constant external input, and the second term is the recurrent input as a function of the average firing rate *r* of the entire network. Both terms depend on the average membrane potential *V_m_*.

We make a final assumption that the firing rate of a neuron depends linearly on its input (linear input-output transfer function). We take this linear dependence to be *r*_out_ = Δ*V_inp_/θ*, where θ = *V_T_ – V_reset_* is the difference between the reset voltage *V*_reset_ and the threshold voltage *V_T_*. Eq. 23 can be rewritten as a self-consistent mean-field equation, given by

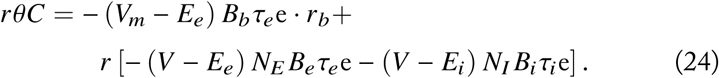

By defining the total baseline input as *S_b_* =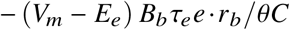 and the total recurrent weight as w = [– 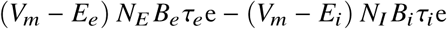] */θC*, we obtain *r* = *S_b_* + *w · r* and therefore *r* = *S_b_*/ (1 − *w*). The stability of the linearized system can be ensured by constraining the total recurrent weight by *w* < 1. For the full network, this provides a condition for stability, given by

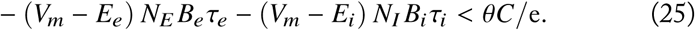

Note that as the left hand side of Eq. 25 depends on the average membrane potential *V_m_* of the network, the condition can be evaluated at different “operating points”. The stability of the excitatory subnetwork, in the absence of the inhibitory subnetwork, can be examined by setting the recurrent inhibitory contribution to zero in Eq 25. This provides a constraint that ensures the network requires inhibitory feedback for stability, given by – (*V_m_* – *E_e_*) *N_E_B_e_τ_e_* ≥ *θC*/e; we therefore obtain the constraint

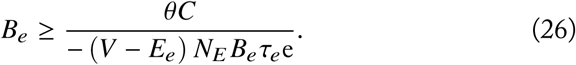

This constraint is plotted as the vertical line denoting the boundary between the ISN and non-ISN regimes in Fig. 5d.

### Experimental Design and Statistical Analysis

No statistical testing was performed. Models and simulations to reproduce all results in this manuscript are available from FigShare DOI 10.6084/m9.figshare.4823212 (https://figshare.com/s/ac309d705ffd3d961fde).

